# The Clathrin adaptor AP-1 and the Rab-stabilizing chaperone Stratum act in two parallel pathways to control the activation of the Notch pathway in *Drosophila*

**DOI:** 10.1101/645580

**Authors:** Karen Bellec, Isabelle Gicquel, Roland Le Borgne

## Abstract

*Drosophila* sensory organ precursors divide asymmetrically to generate pIIa/pIIb cells whose identity relies on the differential activation of Notch during cytokinesis. While Notch is present apically and basally relative to the midbody at the pIIa-pIIb interface, only the basal pool of Notch is reported to contribute to Notch activation in the pIIa cell. Such proper intra-lineage signalling therefore requires appropriate apico-basal targeting of Notch, its ligand Delta and its trafficking partner Sanpodo. We previously reported that AP-1 and Stratum regulate the intracellular trafficking of Notch and Sanpodo from the *trans*-Golgi network to basolateral membrane. Loss of AP-1 or of Stratum caused mild Notch phenotype. Here, we report that the concomitant loss of AP-1 and Stratum result in the stabilization of the apical pool of Notch, Delta and Spdo, the loss of the basal pool of Notch at the pIIa-pIIb interface, and is associated with activation of Notch in the two SOP daughters. We propose that AP-1 and Stratum control two parallel pathways towards plasma membrane and that Notch intra-lineage signalling could also occur at the apical pIIa-pIIb interface.

## Results and Discussion

Cell-cell signalling by the evolutionarily conserved Notch receptor promotes cell fate acquisition in a large variety of developmental processes in metazoans (Artavanis-Tsakonas et al., 1999; Bray, 1998; Kopan and Ilagan, 2009). In most of cases, Notch receptor is activated by transmembrane ligands present at the plasma membrane of adjacent cells. Following binding to Notch, endocytosis of the ligand induces pulling forces driving a change in the conformation of the Notch extracellular domain thereby unmasking the S2 cleavage site of Notch (Gordon et al., 2015; Langridge and Struhl, 2017; Meloty-Kapella et al., 2012; Seo et al., 2016; Shergill et al., 2012; Wang and Ha, 2013). This regulated cleavage is followed by a constitutive proteolytic cleavage of Notch by the gamma secretase complex (Mumm et al., 2000; Struhl and Adachi, 2000) giving rise to the Notch intracellular domain, a polypeptide that is translocated into the nucleus and that acts as a transcriptional coactivator (Artavanis-Tsakonas et al., 1999; Bray, 1998; Kopan and Ilagan, 2009). Because proteolytic activation of the Notch receptor is by definition irreversible, Notch activation needs to be tightly controlled in time and in space. The model system of asymmetric cell division of the sensory organ precursors (SOPs) in the pupal notum of *Drosophila* has been instrumental to identify the site of Notch activation at the cell surface. SOPs are polarized epithelial cells that divide asymmetrically within the plane of the epithelium to generate two daughter cells whose fate depends on the differential activation of Notch signalling (Schweisguth, 2015). It relies on the unequal partitioning of the two cell fate determinants Neuralized (Neur) and Numb in the anterior SOP daughter cell (Le Borgne and Schweisguth, 2003; Rhyu et al., 1994). Neur promotes the endocytosis of Delta (Le Borgne and Schweisguth, 2003) while Numb inhibits the recycling of Notch and its cofactor Sanpodo (Spdo) towards the plasma membrane to instead promote their targeting towards late endosomal compartments (Cotton et al., 2013; Couturier et al., 2013; Johnson et al., 2016; Upadhyay et al., 2013). Consequently, the anterior cell adopts the pIIb identity while Notch is selectively activated in the posterior cell that adopts the pIIa fate. Combination of live-imaging and FRAP experiments using GFP-tagged Notch revealed that proteolytic activation of Notch occurs during SOP cytokinesis and that a specific pool of Notch receptors located basal to the midbody is the main contributor to the signalling in the pIIa cell (Trylinski et al., 2017). These data imply a polarized trafficking of Notch, Delta and Spdo towards this specific subcellular location during cytokinesis. We previously reported that the clathrin adaptor complex AP-1 regulates the polarized sorting of Notch and Spdo from the *trans*-Golgi network (TGN) and the recycling endosomes towards the plasma membrane (Benhra et al., 2011). Loss of AP-1 causes stabilization of Notch and Spdo at the adherens junctions following SOP division, a phenotype associated with a mild *Notch* gain-of-function phenotype (GOF). More recently, we reported that Stratum (Strat), a chaperone regulating Rab8 recruitment, controls the exit from the Golgi apparatus and the basolateral targeting of Notch, Delta and Spdo (Bellec et al., 2018). As for AP-1, loss of Strat causes a mild *Notch* GOF phenotype. These data can be interpreted as AP-1 and Strat acting in the same transport pathway and therefore being simple fine-tune regulators of Notch-Delta trafficking and activation. However, as AP-1 and Strat/Rab8 both regulate basolateral trafficking, an alternative explanation could be that AP-1 and Strat function in two parallel pathways (Fig.1A-A’). In this scenario, loss of one of the two components could be compensated by the other. Accordingly, the concomitant loss of AP-1 and Strat would exhibit a stronger phenotype.

**Figure 1:**
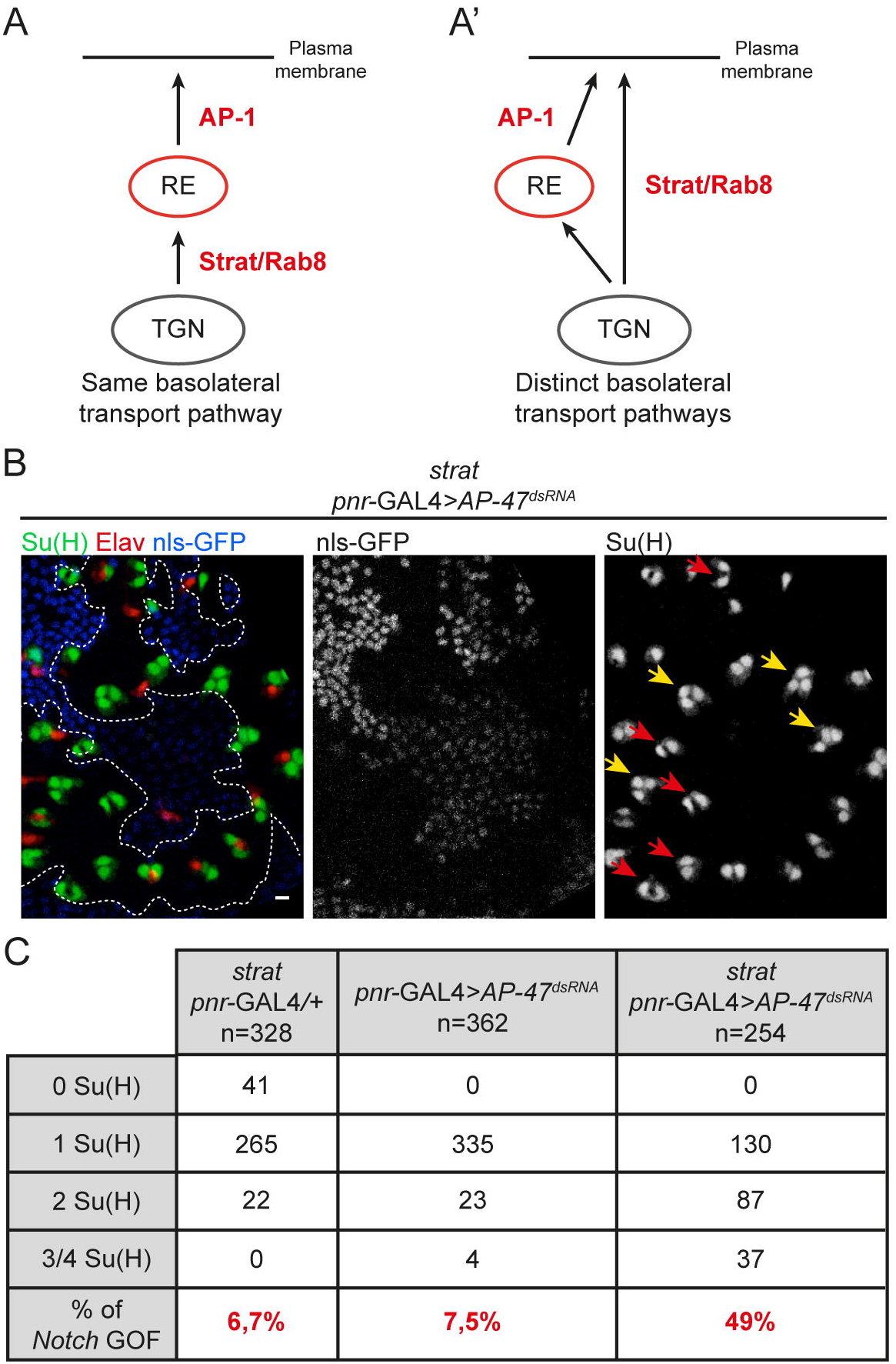
Loss of Strat and AP-1 causes *Notch* gain-of-function phenotype within SO lineage. **A-A’**Schematic representations of the involvement of AP-1 and Strat in the same (**A**) or in distinct (**A’**) basolateral transport pathway. **B.** Projection of confocal sections of a pupal notum at 24h APF. *AP-47*^*dsRNA*^ is expressed using the *pnr*-GAL4 driver and *strat* mutant cells were identified by the absence of the nuclear marker nls-GFP. Sockets were identified with Su(H) (anti-Su(H), green) and neurons with Elav (anti-Elav, red). Scale bar is 5 μm. **C.** Percentage of transformed organs at 24h APF in *strat* mutant, *pnr*-GAL4> + (n=328), in *pnr*-GAL4>*AP-47*^*dsRNA*^ (n=362) and in *strat* mutant, *pnr*-GAL4>*AP-47*^*dsRNA*^ (n=254). Yellow and red arrows show transformed SOs containing 3 to 4 Su(H) positive cells or 2 Su(H) positives cells without neuron, respectively.

### Simultaneous loss of AP-1 and Strat causes a penetrant *Notch* GOF phenotype

To test this prediction, we induced clones of cells homozygote mutant for a null mutation of Strat (Strat KO CRISPR, (Bellec et al., 2018) in which the µ subunit of the AP-1 complex (also known as AP-47) was silenced using the previously described dsRNA (Benhra et al., 2011). Silencing of *AP-47* prevents the assembly of functional AP-1 complexes and gives similar *Notch* GOF phenotypes as two independent *AP-47* null mutant alleles (*AP-47*^*SHE11*^ and *AP-47* deletion by CRISPR ((Benhra et al., 2011) and data not shown)). Silencing of *AP-47* is hereafter referred to as loss of AP-1. In wild-type sensory organ (SO), the pIIb cell divides two times to generate the internal cells among which is one neuron, labelled using an anti-Elav. The pIIa cell divides once to generate the external cells among which is one socket cell, labelled with Suppressor of Hairless (Su(H)). In agreement with previous studies, *Notch* GOF phenotypes corresponding to an excess of socket cells were observed in 6,7% and 7,5% of SO mutant for *strat* or depleted of AP-1, respectively ((Bellec et al., 2018; Benhra et al., 2011), Fig.1C). However simultaneous loss of AP-1 and Strat led to 49% of transformed SOs (Fig.1B-C). Among them, 34% of SOs were transformed at the SOP division, i.e. SOP divided to generate two pIIa cells (n= 87/254, Fig.1C and data not shown). These lineage data indicate that AP-1 and Strat function in two distinct and complementary pathways to regulate Notch activation.

### Numb is correctly partitioned upon loss of AP-1 and Strat

Because AP-1 interacts with Numb and loss of Numb exhibits a *Notch* GOF, we wondered if the phenotype observed upon the loss of AP-1 and Strat would result from the loss of Numb function. We previously reported that the unequal partitioning of Numb is unaffected by the loss of AP-1 (Benhra et al., 2011) or by the loss of Strat (Bellec et al., 2018). Because unequal segregation of Numb is under the control of polarity genes, we first monitored the overall cell polarity (Gho and Schweisguth, 1998; Kraut et al., 1996; Schober et al., 1999). We noticed that, as in *AP-1* mutant, the cuticle looks thinner and less pigmented than in the control situation (**Fig.S1A-B’**). These phenotypes could be caused by a defective AP-1 dependent transport of the Menkes Copper transporter ATP7a that regulates cuticle pigmentation (Holloway et al., 2013; Norgate et al., 2006) and by a defective apical secretion of cuticle components that may rely on AP-1 function as does the apical glue granule secretion in salivary glands (Burgess et al., 2011). Despite these cuticle defects, the localization of the junctional markers DE-Cadherin and Coracle, as well as that of the polarity determinant Par3, were unaffected by the loss of AP-1 and Strat indicating that the overall cell polarity is unaffected (**Fig.S1C-D**). We next monitored the localization of Numb during the division of *strat* mutant SOPs depleted of AP-1. Anti-Numb staining showed that Numb localizes asymmetrically during prometaphase in control and in absence of Strat and AP-1 (Fig.2A-B) and live-imaging using a Numb::GFP^crispr^ (Bellec et al., 2018) revealed that Numb is unequally partitioned in the anterior SOP daughter cell, as seen in the control situation (Fig.2C-D). Thus, the *Notch* GOF phenotype cannot be explained by a defective Numb unequal partitioning during SOP division and this rather suggests that it could be due to a defect in the polarized trafficking of Notch signalling actors.

**Figure 2:**
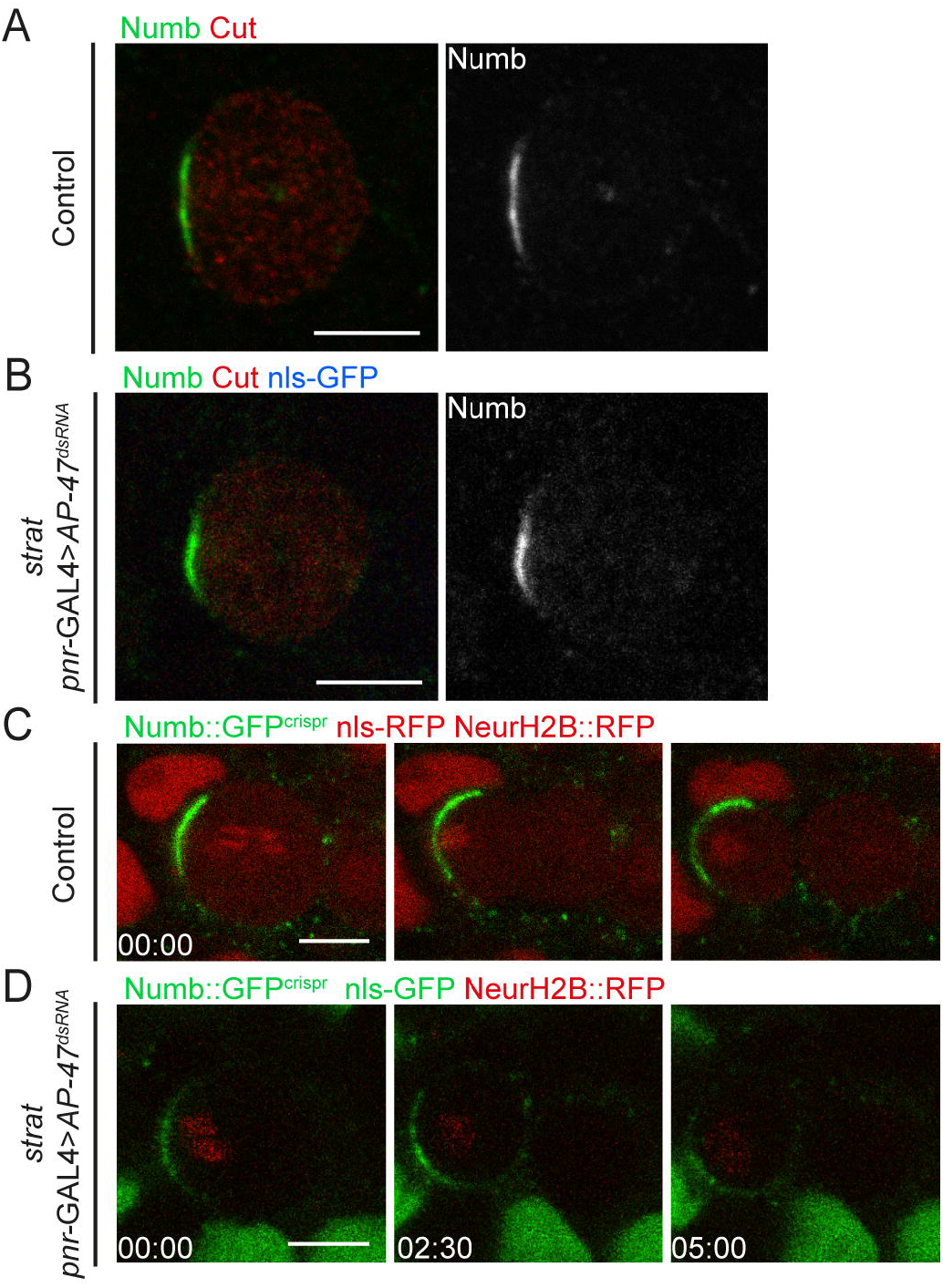
Loss of Strat and AP-1 does not affect the localization of Numb. **A-B.** Localization of Numb (anti-Numb, green) in wild-type (**A**, n=11 SOPs in prophase/prometaphase and n=11 dividing SOPs) and in *strat* SOP expressing *pnr*-GAL4>*AP-47*^*dsRNA*^ (**B**, n=9 SOPs in prophase/prometaphase and n=7 dividing SOPs). SOPs and SOPs daughter cells were identified with Cut (anti-Cut, red). **C-D.** Time-lapse imaging of Numb::GFP^crispr^ (green) in dividing wild-type SOP expressing Histone2B::RFP expressed under the neur promoter (red; **C**, n=10) and in dividing *strat* SOP expressing Histone2B::RFP expressed under the *neur* promoter (red) and *pnr*-GAL4>*AP-47*^*dsRNA*^ (**D**, n=15). Time is minute:second and the time 00:00 corresponds to the SOP anaphase onset. Scale bar is 5 μm.

### Notch is enriched at the apical pIIa-pIIb interface upon loss of AP-1 and Strat

Elegant work from F. Schweisguth laboratory identified two pools of Notch at the pIIa-pIIb interface, apical and basal to the midbody, and provides the compelling evidence that only the subset of receptors located basal to the midbody contributes to signalling in the pIIa cell. To investigate whether AP-1 and Strat affect the apico-basal distribution of Notch receptors, we monitored the dynamics of Notch::GFP^crispr^ (Bellec et al., 2018) throughout the asymmetric division of the SOP. First of all, in the control situation, we confirmed the presence of two pools of Notch along the apical-basal pIIa-pIIb interface, in agreement with previous studies ((Trylinski et al., 2017), Fig.3A). Notch is transiently detected at the apical pIIa-pIIb interface at ~6-9 min after the anaphase onset with a signal intensity peaking at 15-20 min prior to progressively disappear at ~30 min (Fig.3A-B and **Movie S1**). Basal to the midbody, and in agreement with previous studies (Couturier et al., 2012; Trylinski et al., 2017), we also detected punctate structures positive for Notch, hereafter called lateral clusters, that appear ~6-9 min following the anaphase onset and that persist longer than the apical pool of Notch (up to 45 min after the anaphase onset; Fig.3A). In striking contrast to the control situation, upon loss of Strat and AP-1, the Notch lateral clusters were not detected (Fig.3C). Instead, we found that Notch appears in higher amounts at the apical pIIa-pIIb interface cells ~6-9 minutes following the anaphase onset and persists there for longer periods of time compared to the control (at least 78 min post-anaphase versus ~30 min in the control; Fig.3B-C and **Movie S2**). In addition to the apical accumulation, Notch is also detected in dotted structures restricted to the apical plane of the pIIa-pIIb daughter cells (white arrows, t= 39 min and t=78 min, Fig.3C and **Movie S2**). Since lateral clusters from which Notch is activated are absent upon loss of Strat and AP-1, our results raise the possibility that the *Notch* GOF observed could be due to the enrichment of Notch at the apical pIIa-pIIb interface.

**Figure 3:**
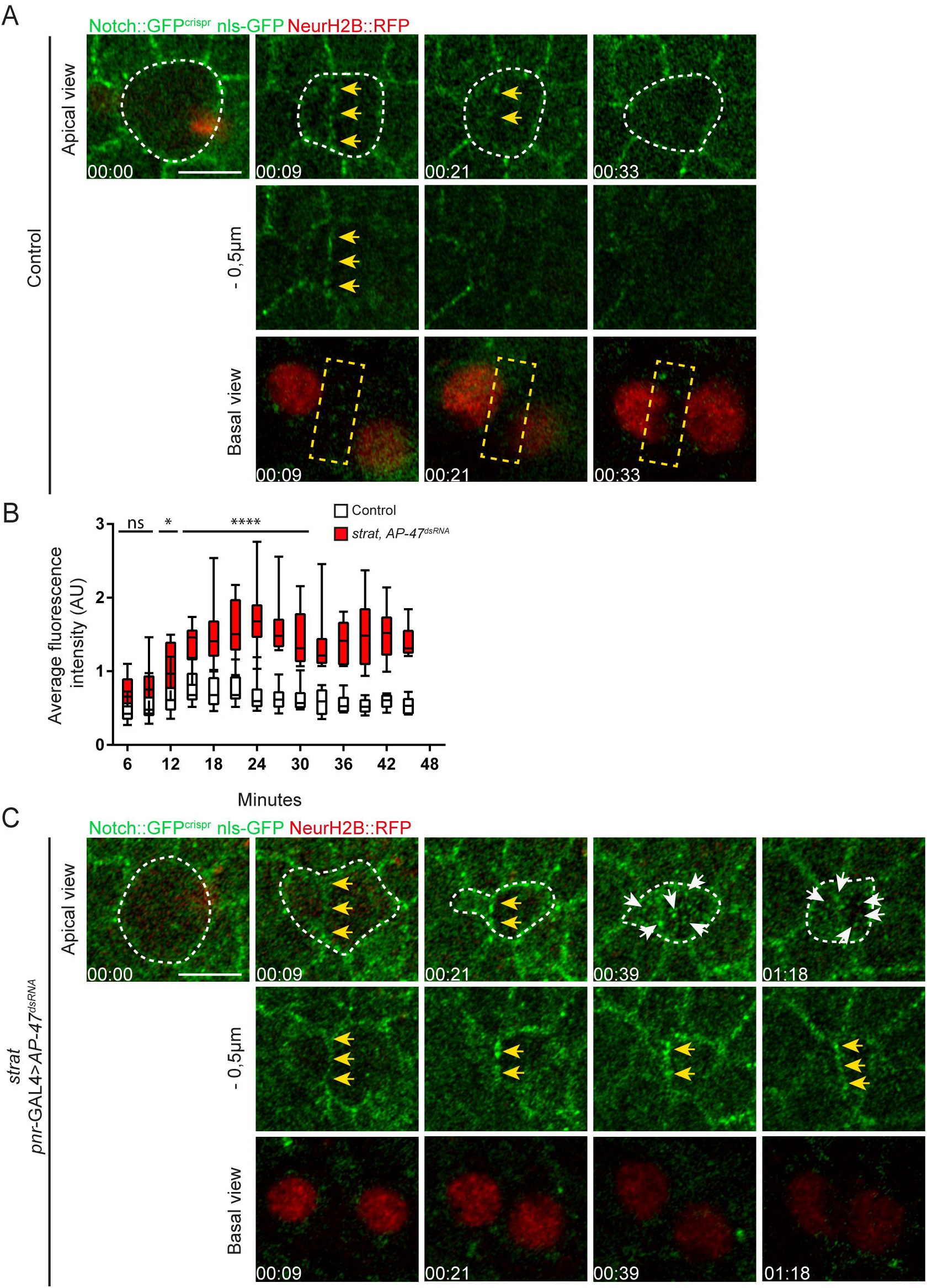
Notch is enriched at the apical pIIa-pIIb interface in the absence of Strat and AP-1. **A.** Time-lapse imaging of Notch::GFP^crispr^ (green) and Histone2B::RFP expressed under the *neur* promoter (red) in dividing wild-type SOP (n=10). **B.** Quantification of the fluorescence intensity of Notch::GFP^crispr^ at the apical pIIa-pIIb interface of wild-type and *strat* SOP daughter cells expressing *pnr*-GAL4>*AP-47*^*dsRNA*^. Boxes extend from the 25^th^ to 75^th^ percentiles and the line in boxes represents the median. The whiskers go down to the smallest value and up to the largest. The statistics were carried out from t=0 to 30min and n=10 for the both conditions (ns≥0.05; \**P*<0.05 and \*\*\*\**P*<0.0001). **C.** Time-lapse imaging of Notch::GFP^crispr^ (green) and Histone2B::RFP expressed under the *neur* promoter (red) in dividing *strat* SOP expressing *pnr*-GAL4>*AP-47*^*dsRNA*^ (n=10/12). Dashed white lines highlight SOP and SOP daughter cells. Yellow arrows point to the enrichment of Notch::GFP^crispr^ at the apical interface between SOP daughter cells and white arrows point to apical compartments positive for Notch::GFP^crispr^. Dashed yellow rectangles highlight compartments positive for Notch::GFP^crispr^ at the basolateral interface between wild-type SOP daughter cells. Basal views are a maximum projection of three confocal slices (Sum projection). Time is in hour:minute and the time 00:00 corresponds to the SOP anaphase onset. Scale bar is 5 μm.

To investigate this possibility, we first tried to directly test the ability of the apical pool of Notch to be cleaved and to be translocated into the nucleus. To this aim, we inserted the green-to-red photoconvertible Dendra2 in the cytoplasmic domain of Notch (Notch::Dendra^crispr^) with the purpose of photoconverting the apical pool of Notch::Dendra^crispr^ and testing its ability to translocate into the nucleus following proteolytic activation. While Dendra2 is detected at the apical plasma membrane upon loss of Strat and AP-1 using an anti-Dendra on fixed specimen (**Fig.S2A**), Notch::Dendra^crispr^ signal is not detectable at the plasma membrane in live specimens (**Fig.S2B**). In time-lapse imaging, Notch::Dendra^crispr^ signal is restricted to intracellular dotted structures, likely endosomal compartments (**Fig.S2B)**. This apparent discrepancy between fixed and living specimen could be due to a longer time of folding or maturation of the newly synthesized Dendra2 compared to the GFP probe, as already reported for other fluorescent probes (Couturier et al., 2014). This result is compatible with the idea that newly synthesized, e.g. not yet folded or matured, Notch::Dendra^crispr^ receptors are targeted to the apical SOP daughter cells interface and are processed from this location upon loss of Strat and AP-1. Because the lack of detection of Notch::Dendra^crispr^ at the plasma membrane prevented us to directly test our hypothesis of apical activation of Notch, we next investigated the localization of Spdo and Delta in SO mutant for Strat and depleted of AP-1.

### Spdo and Delta are distributed with Notch at the apical pIIa-pIIb interface upon loss of Strat and AP-1

The localization of Spdo and Delta was analysed on fixed specimens. As previously described, in the control situation, Spdo is faintly detected at the apical pole of SOP daughter cells, and localizes predominantly in endosomes in the pIIb cell, while in the pIIa cell, Spdo distributes not only in endosomes but also at the basolateral plasma membrane (Fig.4A; (Cotton et al., 2013; Couturier et al., 2013; Hutterer and Knoblich, 2005; Langevin et al., 2005). Upon loss of Strat and AP-1, while Spdo is still detected in dotted intracellular structures in SOP daughter cells, it is no longer detected at the basolateral pIIa-pIIb interface (Fig.4B). Instead, Spdo is enriched at the apical plasma membrane, as observed in single *AP-47^SHE11^* (Benhra et al., 2011) or *strat* mutant background (Bellec et al., 2018; Fig.4B and C) as well as at the apical pIIa-pIIb interface, as observed in most of mutant cases (n=12/14, white arrows, Fig.4B). Similarly to Spdo, and as described previously, Delta is detected into intracellular endocytic structures in pIIa and pIIb cells, at the pIIa-pIIb lateral interface in the control situation as well as the lateral plasma membrane of the pIIa cell. Delta is also weakly detected at the apical pIIa-pIIb interface (Fig. 4D). The loss of Strat and AP-1 leads to an increase in the localization of Delta at the apical pIIa-pIIb interface, together with Notch, in 60% of SOs against 44% in the control situation (Fig.4E and data not shown). Together, our data confirm that, in the control situation, Notch, Delta and Spdo are present at the lateral pIIa-pIIb interface from which Notch activation was reported to take place (Trylinski et al., 2017). Upon loss of Strat and AP-1, sustained levels of Notch/Delta and Spdo at the apical pIIa-pIIb interface are observed, localization at the basolateral interface no longer detected, a phenotype associated with a *Notch* GOF.

**Figure 4:**
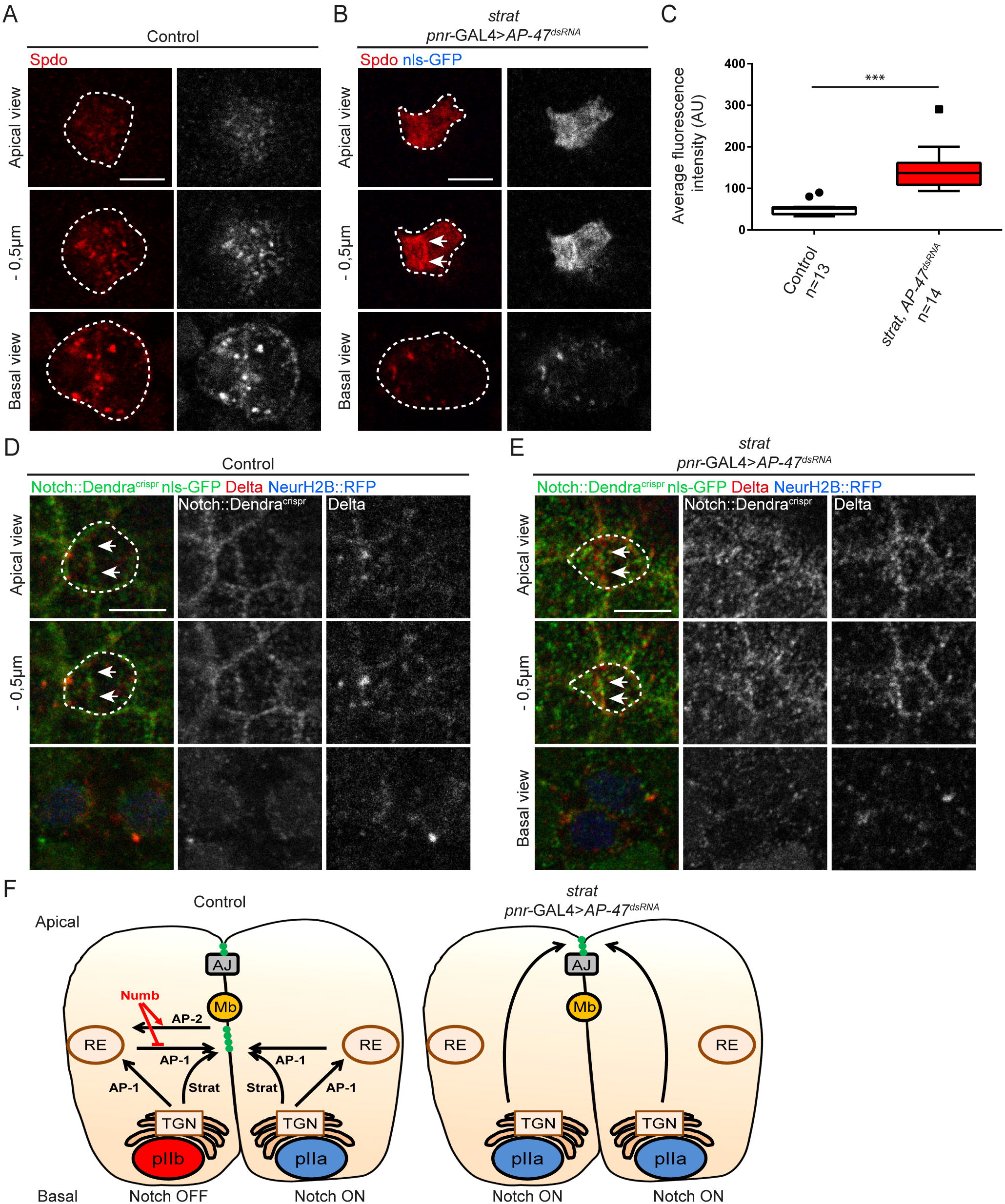
Spdo and Delta are enriched at the apical pIIa-pIIb interface in absence of Strat and AP-1. **A-B.** Localization of Spdo (anti-Spdo, red) in wild-type SOP daughter cells (**A**) and in *strat* SOP daughter cells expressing *pnr*-GAL4>*AP-47*^*dsRNA*^ (**B**). White arrows point to the enrichment of Spdo at the apical interface between SOP daughter cells. **C.** Quantification of the fluorescence intensity of Spdo at the apical pole of wild-type and *strat* SOP daughter cells expressing *pnr*-GAL4>*AP-47*^*dsRNA*^ (\*\*\**P*<0.001). Boxes extend from the 25^th^ to 75^th^ percentiles and the line in boxes represents the median. The circles and the square show the maximum value. **D-E.** Localization of Notch::Dendra^crispr^ (anti-Dendra, green), Histone2B::RFP expressed under the *neur* promoter (blue) and Delta (anti-Delta, red) in wild-type SOP daughter cells (**D**, n=9) and in *strat* SOP daughter cells expressing *pnr*-GAL4>*AP-47*^*dsRNA*^ (**E**, n=20). White arrows point to the enrichment of Notch and Delta at the apical interface between SOP daughter cells. Dashed white lines highlight SOP and SOP daughter cells. Scale bar is 5 μm. **F.** Model of Strat and AP-1 functions in the transport of Notch, Delta and Spdo in wild-type and in *strat* SOP daughter cells expressing *pnr*-GAL4>*AP-47*^*dsRNA*^. Nuclei of cells are in blue (Notch positive) or in red (Notch negative). The traffic of Notch, Delta and Spdo is visualized by black arrows. Notch is represented with green dots. Adherens junctions (AJ) are represented in green, the midbody (Mb) in dark orange, the *trans*-Golgi network (TGN) in light orange and recycling endosomes (RE) are surrounded in brown.

### Concluding remarks

Previous work showed that Strat controls the transport of Notch, Delta and Spdo from the TGN to the basolateral pole (Bellec et al., 2018) and AP-1, from the TGN to the basolateral membrane via recycling endosomes (Benhra et al., 2011; Cotton et al., 2013). Here, we report that AP-1 and Strat act redundantly to control the polarized trafficking of Notch, Delta and Spdo in SOP daughter cells (Fig.4F). Either of these two pathways can be used to ensure the proper localization of Notch, Delta and Spdo, explaining the compensation of one by the other. Upon concomitant loss of Strat and AP-1, Notch, Delta and Spdo are no longer detected in lateral clusters and are enriched at the apical pIIa-pIIb interface, a localization associated with a *Notch* GOF. How can Notch be activated in the absence of lateral clusters, the main contributors of intra-lineage Notch activation (Trylinski et al., 2017)? Although we cannot exclude the possibility that a few, hard to detect lateral clusters could contribute to Notch activation upon loss of AP-1 and Strat, we favour the hypothesis according which Notch activation takes place at the apical pIIa-pIIb interface. The lack of AP-1 prevents Numb to repress the Notch-Spdo recycling normally driving endosomal degradation of Notch-Spdo (Cotton et al., 2013; Couturier et al., 2013; Johnson et al., 2016). Thus, despite its unequal partitioning, Numb is not able to repress Notch activation upon loss of AP-1 and Strat. Nonetheless, although the loss of both of them gives rise to a *Notch* GOF, it does not phenocopy the loss of Numb, where Notch is accumulated in lateral clusters at the pIIa-pIIb interface.

Proteolytic activation of Notch requires the binding of the ligand and its subsequent Neur-mediated endocytosis. While Neur is unequally localized during SOP division (Le Borgne and Schweisguth, 2003), Neur::GFP is detected in both SOP daughter cells, at the apical pIIa-pIIb interface following SOP division (Trylinski et al., 2017). As Delta localizes at the apical pIIa-pIIb interface upon loss of Strat and AP-1, we postulate that Neur can promote endocytosis of Delta in both daughter cells from the apical interface, thereby promoting bidirectional Notch proteolytic activation. If Notch-Delta signalling is primarily taking place at the lateral interface (Trylinski et al., 2017), our study indicates that signalling can also occur apically. The fact that Sec15, a subunit of the exocyst complex, and Arp2/3-WASp (Actin-related protein 2/3-Wiskott-Aldrich syndrome protein) are involved in apical trafficking of Spdo and Delta, respectively, and are required for Notch-Delta signalling is consistent with our proposal of apical activation of Notch (Jafar-Nejad et al., 2005; Rajan et al., 2009). However, the weak contribution of the apical pool of receptors to the Notch activation described in the control situation (Trylinski et al., 2017) suggests that apical activation can only occur when a specific threshold of Notch, Delta and Spdo is reached, such as in *strat* SOs depleted of AP-1. In line with previous reports (Trylinski et al., 2017), our study further illustrates how the apico-basal targeting of Notch, Spdo and Delta along the pIIa-pIIb interface limits signalling between SOP daughter cells at the exit of mitosis. The question of whether this mechanism is conserved awaits further investigation, but given the key role played by Notch in the acquisition of cell fate in vertebrates, it is conceivable that the regulation described in this study may be general.

## Acknowledgements

We thank the Bloomington Stock Center, the Vienna Drosophila RNAi Center and the National Institute of Genetics Fly Stock Center for providing fly stocks. We also thank S. Dutertre and X. Pinson from the Microscopy Rennes Imaging Center-BIOSIT (France). The monoclonal antibodies against Elav, Cut, Cora and DE-Cad were obtained from the Developmental Studies Hybridoma Bank, generated under the auspices of the National Institute of Child Health and Human Development, and maintained by the University of Iowa Department of Biological Sciences. We thank members of R.L.B.’s laboratory for helpful discussions and critical reading of the manuscript.

## Author contributions

Conceptualization: K.B., R.L.B.; Methodology: K.B., I.G., R.L.B.; Formal analysis: K.B., R.L.B.; Investigation: K.B., I.G., R.L.B.; Resources: K.B., I.G.; Writing - original draft: K.B.; Writing - review & editing: K.B., R.L.B.; Visualization: K.B., I.G., R.L.B.; Supervision: R.L.B.; Project administration: R.L.B.; Funding acquisition: K.B., R.L.B.

## Materials and Methods

### *Drosophila* stocks and genetics

*Drosophila melanogaster* stocks were maintained and crossed at 25°C. Mitotic clones were induced using the FLP-FRT technique using the *hs*-FLP and by heat shocking (2×60 min at 37°C) at second and early third instar larvae. *pnr*-GAL4 was used to drive the expression of the *AP-47*^*dsRNA*^. The following stocks were used in this study: *y w hs*-FLP; *Ubi-GFP nls*, FRT40A; *pnr-*GAL4/TM6 *Tb* (Fig.1B,C; 2B,D; 3A-C; 4B-E; **S1B-D; S2; Movies S1 and S2**), *w*; *strat*, FRT40A/CyO; *AP-47*^*dsRNA*^/TM6 *Tb* (stock #24017 from Vienna Drosophila Resource Center, *w*^*1118*^; P(GD14206)v24017/TM3, Fig.1B,C; 2B; 3B,C; 4B,C,E; **S1B-D; S2 and Movie S2**); *y*, *hs*-FLP; *Ubi-GFP nls*, FRT40A (Fig.1C), *w*; *strat*, FRT40A/CyO (Fig.1C); Cad::GFP/+; *pnr-*GAL4/+ (Fig.2A); +/CyO::CFP; *pnr*-GAL4/+ (Fig.2A); *y*, *hs*-FLP; *Ubi-RFP nls*, FRT40A (Fig.2C), *w*; *Numb*::GFP^crispr^, FRT40A/CyO (Fig.2C), *w hs*-FLP *neur-H2B*::RFP;; *Sb*/TM6 *Tb* (Fig.2D), *w*; *strat, Numb*::GFP^crispr^, FRT40A/CyO; *AP-47*^*dsRNA*^/TM6 *Tb* (Fig.2D), *Notch::GFP*^*crispr*^, *neur-H2B*::RFP; *Dr*/TM6 *Tb* (Fig.3A-C; **Movies S1 and S2**), *w*^*1118*^; P(ry[+t7.2]=neoFRT}40A P(w[+mW.hs]=FRT(w[hs]))G13/CyO (stock #8217 from Bloomington, Fig.3A,B; 4D; and **Movie S1**), W1118 (Fig.4A,C), *Notch::Dendra*^*crispr*^, *neur-H2B*::RFP; *Dr*/TM6 *Tb* (Fig. 4D-E; **S1B-B’; and S2**) and *Notch::Dendra*^*crispr*^, *neur-H2B*::RFP; +/+ (**Fig.S1A-A’**).

### Immunofluorescence and antibodies

Pupae were aged for 16.5 to 18.5 h after puparium formation (APF) for SOPs and SOPs daughter cells analysis and were aged for 24 h to 28 h APF for lineage analysis. Pupae were dissected in 1x phosphate-buffered saline (1x PBS) and then fixed for 15 min in 4% paraformaldehyde at room temperature. Dissection and staining conditions were essentially as previously described (Le Borgne and Schweisguth, 2003). Primary antibodies used were rat anti-Elav (7E10, Developmental Studies Hybridoma Bank (DSHB), 1:200), goat anti-Su(H) (sc15813, Santa Cruz, 1:500), mouse anti-Cut (2B10, DSHB, 1:500), goat anti-Numb (SC23579, Santa Cruz, 1:200), rabbit anti-Spdo (a kind gift from J. Skeath, 1:2000) (O’Connor-Giles and Skeath, 2003), mouse anti-DECD (C594.9B, DSHB, 1:200), mouse anti-Cora (C615.16, DSHB, 1:500), rat anti-DE-Cad (DCAD2, DSHB, 1:500), rabbit anti-Par3 (a kind gift from A. Wodarz, 1:1000) (Shahab et al., 2015), mouse anti-HRS (27-4, DSHB, 1:100), rabbit anti-Syntaxin16 (ab32340, Abcam, 1:1000) and rabbit anti-Dendra (Antibodies-online.com, ABIN361314, 1:1000). Cy2-, Cy3- and Cy5-coupled secondary antibodies (1:400) were from Jackson’s Laboratories.

### Generation of Notch::Dendra^crispr^

The Notch::Dendra^crispr^ construct was generated using the CRISPR/Cas9 method as previously described (Gratz et al., 2013a; Gratz et al., 2013b). As for the Notch::GFP^crispr^ construct, the following gRNAs were used: 5′-AACTTGAA|TGGATTGAACCC GGG-3′ and 5′-CGAACTGG|AGGGTTCTCCTGTTG-3′ to introduce Cas9 cuts in exon 6 (Bellec et al., 2018). The Dendra and the 3xP3-DsRed cassette flanked by GVG linkers and by loxP, respectively, were introduced at the previously described position (NiYFP4) (Couturier et al., 2012). Homology arms 1 and 2 were of 1064 bp and 1263 bp in length, respectively. Injection was performed by Bestgene in the *yw*; attP40(nos-cas9)/Cyo stock. The correct position of the Dendra and the DsRed cassette was verified by PCR and sequencing. The DsRed was then removed by crossings with *if*/Cyo, Cre, *w* stock.

### Imaging

Images of fixed nota were acquired with a Leica SPE confocal microscope and a Zeiss Airyscan microscope. The live imaging of Numb::GFP^crispr^ and Notch::Dendra^crispr^ was acquired with a Leica SPE confocal microscope. The live imaging of Notch::GFP^crispr^ in wild-type SOP and in *strat* SOP expressing *pnr*-GAL4>*AP-47*^*dsRNA*^ was acquired with a Zeiss Airyscan microscope. All images were processed and assembled using ImageJ 1.48 and Adobe Illustrator.

### Quantification of the enrichment of Notch at the apical interface

To quantify the signal of Notch::GFP at the apical interface, we measured the signal in a manually drawn area on sum slices of the two apical planes where the Notch signal at the apical interface is the strongest. We also measured the signal between two epithelial cells within an equivalent drawn area and on the same sum slices. Then, we calculated the following ratio: Average fluorescence intensity at the apical interface between SOP daughter cells/Average fluorescence intensity at the apical interface between epithelial cells. This ratio was calculated for each time point. In absence of AP-1 and Strat, the manually drawn area was minimized to avoid taking into account the apical punctate structures positive for Notch.

### Quantification of the apical enrichment of Spdo

The fluorescence intensity was calculated with ImageJ, on two-cell stages, as previously described (Bellec et al., 2018). Briefly, the average fluorescence intensity was measured in a manually drawn area on sum slices of the two most apical planes, where Spdo is enriched. The background noise was measured in the same way and subtracted from the apical intensity value.

### Statistical analysis

Statistical analyses were carried out using the GraphPad Prism 6.05 software. A two-way ANOVA was performed for the quantification of the enrichment of Notch at the apical pIIa-pIIb interface with a multiple comparison Bonferroni test. For the quantification of the apical enrichment of Spdo, because data do not follow a normal distribution, we performed a Wilcoxon test. Statistical significances are represented as follows: not significant (ns)≥0.05; \**P*<0.05; \*\**P*<0.01; \*\*\**P*<0.001 and \*\*\*\**P*<0.0001.

## Supplemental figures

**Figure S1: Apical basal polarity is maintained upon loss of Strat and AP-1**

**A-B’.** Pictures of Notch::Dendra^crispr^ (**A-A’**) and Notch::Dendra^crispr^, *strat* mutant expressing *pnr*-GAL4>*AP-47*^*dsRNA*^ (**B-B’**) at the pupal and adult stages. Dashed black lines highlight depigmented areas. **C.** Localization of Coracle (anti-Cora, green) and DE-Cadherin (anti-DE-cad, red) in wild-type clone expressing *pnr*-GAL4>*AP-47*^*dsRNA*^ and in *strat* mutant clone expressing *pnr*-GAL4>*AP-47*^*dsRNA*^. **D.** Localization of Par3 (anti-Par3, green) in wild-type clone expressing *pnr*-GAL4>*AP-47*^*dsRNA*^ and in *strat* mutant clone expressing *pnr*-GAL4>*AP-47*^*dsRNA*^. Dashed yellow lines delineate *strat* mutant clones. Scale bars are 500 µm for pictures and 5 μm for immunostainings.

**Figure S2: Notch::Dendra^crispr^ is only detected in endosomal compartments in live specimen**

**A.** Localization of Notch::Dendra^crispr^ (green) and Dendra (anti-Dendra, red) in a *strat* notum expressing *pnr*-GAL4>*AP-47*^*dsRNA*^. Dashed white lines highlight the *strat* mutant clone. **B.** Time-lapse imaging of Notch::Dendra^crispr^ (green) and Histone2B::RFP expressed under the *neur* promoter (red) in dividing *strat* SOP expressing *pnr*-GAL4>*AP-47*^*dsRNA*^. Time is in minute:second and the time 00:00 corresponds to the SOP anaphase onset. Scale bar is 5 μm.

## Supplemental movies

**Movie S1: Notch dynamic in dividing wild-type SOP**

Apical plan of a time-lapse imaging of Notch::GFP^crispr^ (green) in dividing wild-type SOP (n=10). Time is in minutes and the time 0 corresponds to the SOP anaphase onset.

**Movie S2: Notch dynamic in dividing *strat* SOP expressing *pnr*-GAL4>*AP-47*^*dsRNA*^**

Apical plan (Projection of the 2 most apical slices) of a time-lapse imaging of Notch::GFP^crispr^ (green) in dividing *strat* SOP expressing *pnr*-GAL4>*AP-47*^*dsRNA*^ (n=10). Time is in minutes and the time 0 corresponds to the SOP anaphase onset.

